# Strain-specific early, host-shaped late: the rhizobial transcriptome across symbiosis

**DOI:** 10.64898/2026.11.11.765592

**Authors:** Sebastián Acosta-Jurado, Francisco Fuentes-Romero, Pilar Navarro-Gómez, Paula Ayala-García, Francisco Javier Ollero, José Ignacio Jiménez-Zurdo, Francisco Pérez-Montaño, José María Vinardell, Francisco Javier López-Baena

**Affiliations:** Departamento de Microbiología, Facultad de Biología, Universidad de Sevilla, Avda. Reina Mercedes 6, 41012 Sevilla, Spain; Structure, Dynamics and Function of Rhizobacterial Genomes (RhizoRNA Lab), Estación Experimental del Zaidín, Consejo Superior de Investigaciones Científicas (CSIC), Granada, Spain

**Keywords:** rhizobia, bacteroid, transcriptomics, nodule type, host specificity, type III secretion, Nod factors, *Sinorhizobium fredii*, *Rhizobium tropici*

## Abstract

Early flavonoid signalling and mature bacteroid differentiation represent the two ends of the rhizobial symbiotic trajectory. Here, we report strand-specific RNA-seq profiles of mature bacteroids compared against free-living cells and early flavonoid responses in two strains: *Sinorhizobium fredii* HH103 from *Glycine max*, *Glycyrrhiza uralensis*, and *Lotus burttii*, and *Rhizobium tropici* CIAT 899 from *Phaseolus vulgaris* and *L. burttii*, the sole legume nodulated by both strains. A host-common core dominates these transcriptomes, with 1,171 of 2,855 responding genes shared across all three HH103 hosts and 1,664 of 2,703 across both CIAT 899 hosts. What distinguishes the remainder is host identity rather than nodule type, as HH103 bacteroids from the indeterminate-nodule host *G. uralensis* resemble those from *L. burttii* more closely than those from *G. max*. Nitrogen fixation and stress responses are over-represented in every bacteroid, whereas motility is almost entirely repressed. Dismantling of the early programme is selective: in HH103, the type III secretion system is switched off, the Nod factor operons are attenuated but retained, and the *nod*-box-regulated *fixABCX* operon is amplified. CIAT 899, which lacks a type III system, keeps its *nif/fix* gene block inactive until nodule formation, when it engages 26 of its 30 genes. Across genera, 26% of responding orthologue pairs are shared in the bacteroid versus 5% during the flavonoid response: each bacterium negotiates entry in a dialect of its own and converges on a shared end state whose amplitude, rather than content, the host modulates.

**Graphical abstract:** 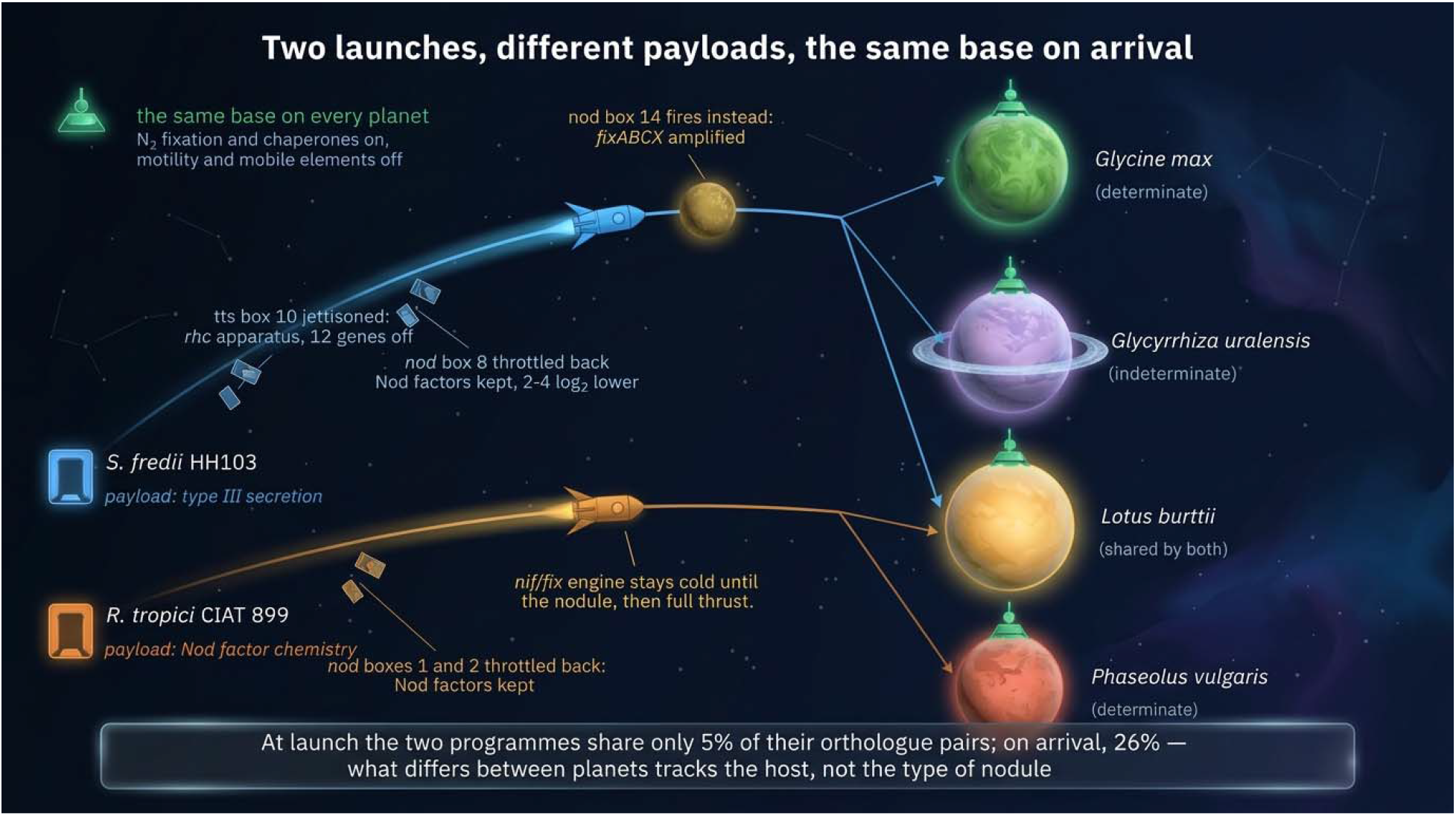

## 1. Introduction

The symbiosis between legumes and rhizobia converts atmospheric nitrogen inside a dedicated root organ, and it is governed by a strictly enforced molecular dialogue in which the plant controls every step of bacterial entry and accommodation (Oldroyd et al., 2011; Oldroyd, 2013). For the bacterium the interaction entails a complete change of life, from free-living soil saprophyte to intracellular, differentiated endosymbiont (Poole et al., 2018). The exchange begins in the rhizosphere: flavonoids exuded by the legume root are perceived by the rhizobial LysR-type regulator NodD, which binds conserved promoter sequences known as *nod* boxes and activates the genes that synthesise Nod factors, the lipochitooligosaccharides that trigger root hair curling and the nodulation programme in the plant (Kobayashi et al., 2004). Nod factors are the universal currency of the interaction, but not the whole message: flavonoid perception also mobilises, depending on the species, surface polysaccharide production, phytohormone synthesis, motility, quorum sensing and protein secretion systems (Jiménez-Guerrero et al., 2018). Which legumes a strain can nodulate is therefore set by that whole combination of molecular keys and not by Nod factors alone (Perret et al., 2000).

Of the accessory determinants, the Type III Secretion System (T3SS) is the best characterised. In rhizobia that possess one, the NodD-dependent regulator TtsI recognises a second class of promoter elements, the *tts* boxes, and switches on the secretion apparatus and the nodulation outer proteins (Nops) delivered through it into the host cytoplasm (López-Baena et al., 2008). These effectors are double-edged: the same protein can promote nodulation of one legume and abort it in another, depending on whether the host recognises it (Staehelin and Krishnan, 2015; Jiménez-Guerrero et al., 2022), and in *Bradyrhizobium elkanii* the T3SS can even trigger nodule formation without Nod factors or their receptors (Okazaki et al., 2013). A symbiotic T3SS is nevertheless not a general feature of rhizobia (Jiménez-Guerrero et al., 2022).

HH103 and CIAT 899 are both broad-host-range symbionts that reach that breadth by different means, and the contrast is the reason for pairing them. *S. fredii* HH103, whose 7.25 Mb genome is distributed over one chromosome and six plasmids and is the structurally most complex sinorhizobial genome described (Vinardell et al., 2015; Fuentes-Romero et al., 2025), nodulates legumes spanning determinate and indeterminate nodule types, and a substantial part of that range is set by its T3SS: 17 *rhc* genes encode the secretion apparatus and distinct Nops are delivered through it, most of them encoded on the symbiotic plasmid. These effectors act as symbiotic factors on some hosts and as asymbiotic ones on others, widening the range with *Glycine max* and *Glycyrrhiza uralensis* while restricting it with *Erythrina variegata* or *Lotus japonicus* Gifu (López-Baena et al., 2008; Jiménez-Guerrero et al., 2022).

*R. tropici* CIAT 899, an acid- and salt-tolerant strain used as a commercial inoculant for common bean, has no T3SS at all: its genome encodes secreted adhesins and type I and type V secretion systems but no type III apparatus, and no *rhc* or *nop* genes (Ormeño-Orrillo et al., 2012). It negotiates entry through amplified signal chemistry instead. Its symbiotic plasmid carries five *nodD* regulatory genes and three *nodA* genes, the largest complement described in a rhizobium, which underlie the most complex regulation of nodulation genes known and an exceptionally diverse family of Nod factors (Ormeño-Orrillo et al., 2012; del Cerro et al., 2015). Genetic dissection of that redundancy has shown that only NodD1 and NodD2 contribute to symbiosis, and that either *nodA1* or the flavonoid-independent *nodA3* suffices for nodulation (del Cerro et al., 2019a; Ayala-García et al., 2022). Its most distinctive property is the abundant production of Nod factors under non-inducing conditions, and above all under osmotic stress: ionic stress with NaCl and non-ionic stress with mannitol both switch on the *nod* genes in the complete absence of flavonoids, with NodD2 as the regulator responsible (del Cerro et al., 2017, 2019b), while acidity elicits a distinct profile of its own (Morón et al., 2005; Estévez et al., 2009; Guasch-Vidal et al., 2013).

Both strains were profiled at that early, free-living stage ten years ago. In HH103, genistein elicited 106 differentially expressed genes, 101 of which were sorted into three classes according to the regulatory element upstream: genes preceded by a *nod* box, genes preceded by a *tts* box, and genes belonging to neither category, whose regulation could not be explained (Pérez-Montaño et al., 2016a). The equivalent experiment in CIAT 899, with apigenin and with salt, returned only 19 genes responding to the flavonoid, 15 of them induced and located on the symbiotic plasmid, with the two inducers converging on a largely overlapping set of symbiotic genes (Pérez-Montaño et al., 2016b). Both belong to a body of work covering the flavonoid response of *Bradyrhizobium japonicum* USDA 110, *Rhizobium leguminosarum* bv. viciae 3841, *Sinorhizobium meliloti* 1021, *R. tropici* CIAT 899 and *S. fredii* HH103, compiled by Jiménez-Guerrero et al. (2018), whose central conclusion is that each species brings its own arsenal of molecular weapons to accompany the shared Nod factor signal.

This early work was produced with non-strand-specific RNA-seq, a choice whose cost is particularly acute in bacteria. Without strand information, a read falling where transcription occurs on both strands cannot be assigned to the gene that produced it, and bacterial genomes are compact, with densely packed operons, overlapping ends and pervasive antisense transcription that amounts to a regulatory layer of its own (Georg and Hess, 2011). The concern is not hypothetical in rhizobia: in *S. meliloti*, antisense transcription covers a large share of protein-coding genes (Jiménez-Zurdo and Robledo, 2015) and is biased towards the nodulation and nitrogen fixation genes of the symbiotic megaplasmid (Robledo et al., 2015), precisely the genes such experiments set out to quantify. Protocols preserving the strand of origin were benchmarked and standardised well before those studies were performed (Levin et al., 2010). The consequence of omitting them is not a uniform loss of sensitivity: strongly induced genes remain detectable because their own signal dominates, whereas genes of modest amplitude are the ones whose quantification is distorted. A regulon defined on non-strand-specific data is therefore expected to be reliable at its core and unreliable at its periphery.

The second and larger gap is developmental. Everything known transcriptionally about HH103 and CIAT 899 concerns free-living cells responding to a purified flavonoid, a condition that captures the first hours of a dialogue lasting weeks. No transcriptome of either strain has been reported for the bacteroid, the differentiated, nitrogen-fixing form adopted inside the symbiosome, even though it is the stage at which the symbiosis delivers its function. A description covering the same genome from the earliest recognition events through to the mature bacteroid, and across several hosts rather than one, would turn two disconnected snapshots into a continuous account of the interaction, and would allow the central questions to be put directly: what the bacterium carries forward from recognition into fixation, what it discards, and the extent of each depends on the host plant it colonizes.

The bacteroid is also not a single state, because the organ that houses it is not a single structure. Determinate nodules, as in soybean, *Lotus* and common bean, lose their meristem early and remain spherical with a broadly synchronous bacterial population, whereas indeterminate nodules, as in *Medicago*, pea and *Glycyrrhiza*, keep a persistent apical meristem and elongate, holding a developmental gradient from meristem to senescence within one organ (Oldroyd et al., 2011). Nodule type is often assumed to dictate the fate of the symbiont, because legumes of the inverted repeat-lacking clade (IRLC) impose terminal differentiation through nodule-specific cysteine-rich (NCR) peptides (Van de Velde et al., 2010), the severity of the effect scaling with how many NCRs the host encodes (Montiel et al., 2017). The two properties are separable, however, and the hosts used here separate them: *G. uralensis* is an IRLC legume with few NCRs, and HH103 bacteroids from its nodules keep the DNA content, size and viability of free-living cells (Crespo-Rivas et al., 2016).

Bacteroid transcriptomes are well established in other systems. *S. meliloti* has been profiled at sub-nodule resolution in *Medicago truncatula* by laser-capture microdissection coupled to RNA sequencing (Roux et al., 2014), and three studies have compared bacteroids of a single genome between nodule types: *Sinorhizobium* sp. NGR234 in determinate *Vigna unguiculata* and indeterminate *Leucaena leucocephala* (Li et al., 2013), *R. leguminosarum* in determinate *Phaseolus* and indeterminate pea using isogenic strains differing in their symbiosis plasmid (Green et al., 2019), and *Bradyrhizobium arachidis* in determinate peanut and indeterminate *Sophora flavescens* (Chen et al., 2023). Others have compared hosts within a single nodule type, for *R. leguminosarum* in pea and *Vicia cracca* (Karunakaran et al., 2009) and for *Bradyrhizobium japonicum* in soybean and cowpea (Koch et al., 2010). Pronounced differences between nodule types are reported (Green et al., 2019), yet at the level of morphology a survey of 40 Papilionoideae species found no clear correlation between nodule type and bacteroid form (Oono et al., 2010).

To our knowledge, what has not been done in any system is to follow the same two bacteria along that whole trajectory. The published studies are either free-living or symbiotic, and those that compare hosts use a single determinate and a single indeterminate legume, which confounds nodule type with host identity. Here we report strand-specific RNA-seq of HH103 bacteroids from soybean (*Glycine max*), *Lotus burttii* and *Glycyrrhiza uralensis*, and of CIAT 899 bacteroids from common bean (*Phaseolus vulgaris*) and *L. burttii*, each contrasted against free-living cells of the same strain, together with a re-examination of the genistein and apigenin responses. Three features of the design address the confounds above: each strain is sampled on its own natural host as well as on hosts of its extended range; both nodule types occur within one bacterial genotype, *G. max* and *L. burttii* being determinate and *G. uralensis* indeterminate, while the differentiation state stays constant across the three; and one legume, *L. burttii*, is shared by the two bacteria, making the comparison between genera a controlled one. Confining the nodule-type contrast to HH103 is deliberate: CIAT 899 does nodulate indeterminate hosts such as *Leucaena leucocephala*, but whether its bacteroids differentiate terminally in them has never been determined, so any difference found would confound nodule architecture with differentiation fate. The re-examination of the 2016 conditions first establishes what the earlier, non-strand-specific datasets recovered and what they missed; the bacteroid data then address how much of the programme is shared among hosts and how much is host-specific, whether nodule type organises that variation, and how far the programme is conserved between two rhizobia that share a legume but not a genus.

## 2. Materials and methods

### 2.1 Bacterial strains and growth conditions

*Sinorhizobium fredii* HH103 and *Rhizobium tropici* CIAT 899 were grown at 28 °C in Minimal Medium (MM; Robertsen et al., 1981). For the free-living conditions, three independent 25 mL cultures were set up per condition and supplemented with the corresponding *nod* gene-inducing flavonoid, genistein for HH103 and apigenin for CIAT 899, dissolved in ethanol to a final concentration of 3.7 µM; control cultures received an equivalent volume of ethanol. Cells were harvested by centrifugation 48 h after the addition of the flavonoid, and the pellets were stored at -80 °C until RNA extraction.

### 2.2 Plant material, inoculation and nodule harvest

HH103 was inoculated onto soybean (*G. max* cv. Williams), *L. burttii* and *G. uralensis*, and CIAT 899 onto common bean (*P. vulgaris* cv. BBL) and *L. burttii*. Nodulation assays followed the procedures already established in this laboratory for each legume. For soybean, common bean and *L. burttii* they were performed as described by Hidalgo et al. (2010) and Acosta-Jurado et al. (2016), common bean following the same procedure as soybean: surface-sterilised seeds were pregerminated and transferred to Leonard jar assemblies, two plants per jar for soybean and common bean and four for *Lotus*, and each plant was inoculated with approximately 5 x 10^8^ bacteria for soybean and common bean and 10^8^ for *Lotus*. For *G. uralensis* they followed Mergaert et al. (2003) and Crespo-Rivas et al. (2009): germinated seeds were transferred to Leonard jars containing sterilised vermiculite supplemented with nitrogen-free Fahraeus nutrient solution (Vincent, 1970) and each plant was inoculated with approximately 10^8^ bacteria. In every case plants were grown in a chamber under a 16 h photoperiod at 25 °C in the light and 18 °C in the dark.

Nodules were collected approximately six to seven weeks after inoculation for *G. max*, *L. burttii* and *P. vulgaris*, and about two months after inoculation for the slower-nodulating *G. uralensis*. Three biological replicates were analysed per strain-host combination. Each replicate came from an independent nodulation assay, so that the three are biologically independent and not aliquots of a common harvest; within each assay the nodules of the inoculated plants were collected and pooled, giving of the order of 40 nodules per replicate, which for soybean corresponds to approximately five to ten plants.

### 2.3 Bacteroid isolation

Bacteroids were extracted as described by Finan et al. (1991), following the procedure applied to HH103 by Crespo-Rivas et al. (2016). Nodules were surface-sterilised for 1 min in 100% bleach and washed three times in water. They were then crushed in MMS buffer (40 mM 3-(4-morpholino)-propane sulfonic acid, 20 mM KOH, 2 mM MgSO4, 0.3 M sucrose, pH 7.0) and the liquid containing the bacteroids was removed with a pipette. Large particles were eliminated by passage through a 40 µm cell strainer cap (BD Falcon, BD Biosciences, San Jose, CA, USA). Samples were centrifuged once at 70 × g for 5 min and subsequently twice at 2,200 × g for 5 min, resuspending the resulting pellet fraction in MMS buffer. No further plant-RNA depletion step was applied: the separation of bacteroids from plant material relies on this differential centrifugation, and residual host transcripts are removed in silico at the mapping stage, since reads are aligned against the rhizobial genome only.

### 2.4 RNA extraction and purification

Total RNA was extracted from free-living cells and from isolated bacteroids by the same procedure, using TRI Reagent (Invitrogen, USA) under strict RNase-free conditions. Frozen pellets stored at -80 °C were thawed on ice and TRI Reagent was added at a ratio of 2 mL per 25 mL of culture pellet. Mixtures were incubated at 65 °C for 20 min and centrifuged at 13,000 × g and 4 °C for 10 min. Phase separation was initiated by adding 0.1 volumes of 1-bromo-3-chloropropane to the supernatant, vortexing for 30 s, equilibrating for 5 min at room temperature and centrifuging at 18,000 × g and 4 °C for 15 min. The upper aqueous phase was transferred to a clean tube, avoiding the interphase, and the RNA was precipitated with one volume of isopropanol for 10 min at room temperature (13,000 × g, 4 °C, 20 min), washed with 1 mL of ice-cold 70% ethanol prepared in diethyl pyrocarbonate (DEPC)-treated water (13,000 × g, 4 °C, 5 min), dried at 55-65 °C and resuspended in 50 µL of pre-heated DEPC-treated water.

Residual genomic DNA was digested by incubating 100 µL of RNA with 5 µL of DNase (Roche, Switzerland), 5 µL of RNase inhibitor and 10 µL of 10× buffer at 37 °C for 1 h. The reaction was diluted with 100 µL of DEPC-treated water and purified by two sequential extractions with one volume of acid phenol:chloroform:isoamyl alcohol (25:24:1, v/v/v), centrifuging at 18,000 × g and 4 °C for 10 min. The final aqueous phase was precipitated overnight at -20 °C with three volumes of ice-cold absolute ethanol and 0.1 volumes of 3 M sodium acetate (pH 5.2), pelleted (18,000 × g, 4 °C, 30 min), washed with ice-cold 70% ethanol, dried at 60 °C for 10 min and resuspended in 50 µL of DEPC-treated water. Samples were finally cleaned up with the RNeasy Plus Mini Kit (Qiagen) following the manufacturer’s instructions, including the gDNA Eliminator columns, and eluted in 30-50 µL of RNase-free water. RNA quantity, purity and integrity were assessed with a Nanodrop One spectrophotometer, a Qubit fluorometer and a 2100 Bioanalyzer.

### 2.5 Library preparation and sequencing

Library construction and sequencing were performed at the Genomics Unit of the Instituto de Parasitología y Biomedicina “López-Neyra” (IPBLN-CSIC, Granada, Spain), project NGS008/21. Because of the prevalence of small RNA species in the preparations, samples were first subjected to a partial depletion of these molecules by solid-phase reversible immobilisation at 3.5×. Strand-specific libraries were then built from 200 ng of total RNA with the Illumina Stranded Total RNA Prep with Ribo-Zero Plus kit and IDT for Illumina RNA UD Indexes Set B, and their size distribution was verified on a Bioanalyzer High Sensitivity DNA chip. Sequencing was carried out on an Illumina NextSeq platform in Mid Output 150-cycle configuration, paired-end 2 × 72 nt, over two runs that yielded 75.0 and 73.9 Gbp, corresponding to 457.7 and 451.1 million passing-filter reads, with Q30 values of 90.5% and 92.0%.

### 2.6 Read mapping and differential expression

Differential expression was called with DESeq2, which models counts with a negative binomial distribution, normalises libraries by the median of ratios and corrects for multiple testing by the Benjamini-Hochberg procedure. For every contrast, bacteroid or flavonoid-treated samples were compared against free-living control cells of the same strain, with three biological replicates per condition. Genes were considered differentially expressed at a fold change of at least 3 in absolute value and an adjusted *P*<0.005. Reads were aligned to the reference genomes with Rbowtie2 2.2.0, alignments handled with Rsamtools 2.2.3 and samtools 1.15.1, intervals processed with bedtools 2.30.0, and reads assigned to features with Rsubread 2.10.4; expression levels and differential expression were computed with the Bioconductor package DESeq2 1.36.0.

### 2.7 Reference genomes and annotation

Reads were referred, for HH103, to the closed HH103_v32 assembly of the strain (Fuentes-Romero et al., 2025), whose seven replicons are the chromosome (CP183939), pSfHH103a1 (CP183940), pSfHH103a2 (CP183941), pSfHH103b (CP183942), pSfHH103c (CP183943), pSfHH103d/pSym (CP183945) and pSfHH103e (CP183944), and for CIAT 899 to the complete genome of Ormeño-Orrillo et al. (2012), comprising the chromosome (CP004015) and the plasmids pRtrCIAT899a (CP004016), pRtrCIAT899b (CP004017) and pRtrCIAT899c (CP004018). The symbiotic plasmid is pRtrCIAT899b, which carries all five *nodD* and all three *nodA* genes and 47 of the 51 genes whose names begin with *nod*, *nol*, *noe*, *nif* or *fix*; the other four, *noeJ*, *noeK*, *nolR* and *nodN*, are chromosomal, and pRtrCIAT899c, although the largest plasmid, carries none. The HH103 annotation resolves into 7,010 locus tags, 6,947 coding sequences and 63 structural RNA genes (54 tRNA, 9 rRNA); these are the identifiers used in the expression tables, so each carries an annotation record. Two coding tags label two separate segments each, SFHH103_02179, split in two, and the transposase SFHH103_06138, present at two loci on pSfHH103e, which is why the assembly lists 7,012 feature records. The structural RNA genes were kept in the differential-expression tables as supplied and account for 38 of the DEGs reported here. The CIAT 899 assembly has not changed, but its annotation exists in two generations: the original tags RTCIAT899_CH, _PA, _PB and _PC, quoted in Pérez-Montaño et al. (2016b), and the RefSeq reannotation of the same sequences (GCF_000330885.1), renamed RTCIAT899_RS, which is the one used here. Both were reconciled through the old_locus_tag field: 6,140 of the 6,382 genes carry such a link, the other 242 being RefSeq calls absent from the original annotation. A product description is available for 6,324 of those features, and the “Not annotated” block of Table 1 collects the 325 HH103 and 54 CIAT 899 features that lacked one when the functional blocks were built, so that block measures coverage of identifiers rather than of function. Table S7 gives the full annotation of both strains.

**Table 1.**
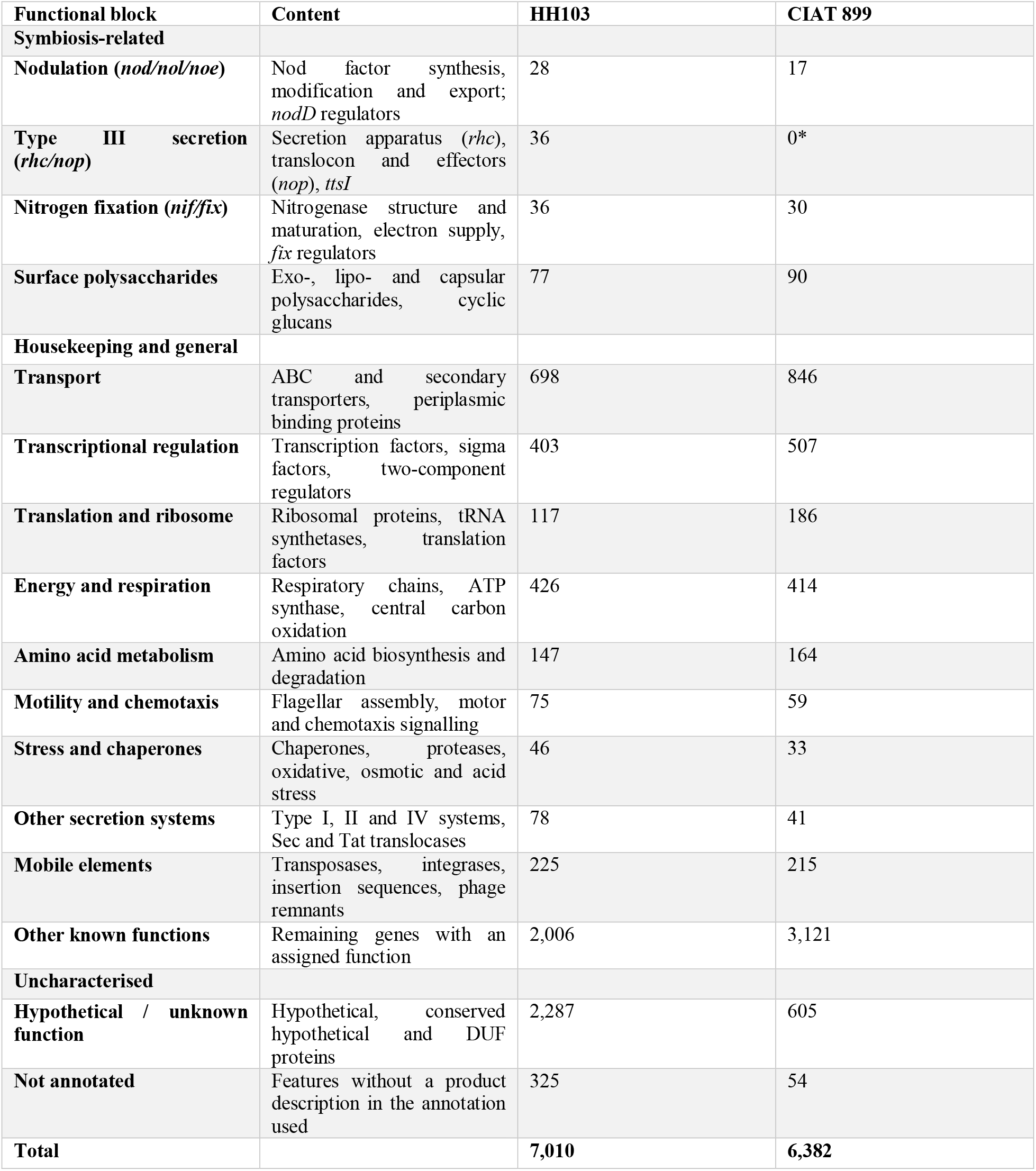
The sixteen functional blocks and their size in each genome. Every protein-coding gene is assigned to exactly one block from its gene name, product and annotation note, with symbiotic categories tested before general ones; the “Content” column states what each block collects, and the two right-hand columns give the number of genes it contains in each genome, which is the denominator of the percentages plotted in Figures 2 and 4. *CIAT 899 encodes no T3SS. Full per-gene assignments are given in Table S7, and the enrichment statistics computed over these blocks in Table S4.

| Functional block | Content | HH103 | CIAT 899 |
|---|---|---|---|
| **Symbiosis-related** |  |  |  |
| **Nodulation (*nod/nol/noe*)** | Nod factor synthesis, modification and export; *nodD* regulators | 28 | 17 |
| **Type III secretion (*rhc/nop*)** | Secretion apparatus (*rhc*), translocon and effectors (*nop*), *ttsI* | 36 | 0* |
| **Nitrogen fixation (*nif/fix*)** | Nitrogenase structure and maturation, electron supply, *fix* regulators | 36 | 30 |
| **Surface polysaccharides** | Exo-, lipo- and capsular polysaccharides, cyclic glucans | 77 | 90 |
| **Housekeeping and general** |  |  |  |
| **Transport** | ABC and secondary transporters, periplasmic binding proteins | 698 | 846 |
| **Transcriptional regulation** | Transcription factors, sigma factors, two-component regulators | 403 | 507 |
| **Translation and ribosome** | Ribosomal proteins, tRNA synthetases, translation factors | 117 | 186 |
| **Energy and respiration** | Respiratory chains, ATP synthase, central carbon oxidation | 426 | 414 |
| **Amino acid metabolism** | Amino acid biosynthesis and degradation | 147 | 164 |
| **Motility and chemotaxis** | Flagellar assembly, motor and chemotaxis signalling | 75 | 59 |
| **Stress and chaperones** | Chaperones, proteases, oxidative, osmotic and acid stress | 46 | 33 |
| **Other secretion systems** | Type I, II and IV systems, Sec and Tat translocases | 78 | 41 |
| **Mobile elements** | Transposases, integrases, insertion sequences, phage remnants | 225 | 215 |
| **Other known functions** | Remaining genes with an assigned function | 2,006 | 3,121 |
| **Uncharacterised** |  |  |  |
| **Hypothetical / unknown function** | Hypothetical, conserved hypothetical and DUF proteins | 2,287 | 605 |
| **Not annotated** | Features without a product description in the annotation used | 325 | 54 |
| **Total** |  | **7,010** | **6,382** |

### 2.8 Analytical methods

Differentially expressed genes (DEGs) were defined across both strains and all conditions using a uniform threshold of a fold change of at least 3 (log_2_≥1.585) and a Benjamini–Hochberg–adjusted P value less than 0.005, restricted to protein-coding features. Functional classification combined two complementary schemes in parallel: a custom 16-category functional block scheme with 100% coverage that assigns each coding gene to a single category via regular expressions over gene name, product, and annotation note under an explicit precedence hierarchy and an organism-independent COG assignment obtained via NCBI Batch CD-Search (best COG hit per protein at an E value less than 0.01). COG assignments covered 10,235 of the 13,392 proteins across both strains (76%), comprising 5,338 of 7,010 proteins in HH103 and 4,897 of 6,382 in CIAT 899, with genes assigned to multiple COGs contributing to each category. Finally, orthologues were identified as reciprocal best BLASTP hits meeting at least 30% amino-acid sequence identity and at least 50% query coverage. Three constraints apply to these analyses. Predicted non-coding RNAs were excluded throughout (6,733 of 13,743 features in HH103 and 14,339 of 20,721 in CIAT 899). One contiguous block of 400 chromosomal CIAT 899 genes (RTCIAT899_RS15570-RS17600) could not be submitted to CD-Search and was removed from both the differentially expressed set and the background of that strain, so the universe of the COG enrichment test is 4,696 genes for CIAT 899 against 5,338 for HH103. In the apigenin contrast 2,538 genes lack an adjusted P value, against 133 in each bacteroid contrast, so the 274 apigenin DEGs derive from a smaller testable universe than the other conditions. No count matrices or replicate-level data were available, so no ordination or re-estimation of dispersion was performed and the differential-expression statistics are taken as provided.

## 3. Results

### 3.1 Strand-specific libraries reproduce the decade-old datasets and extend them

Both strains had been profiled in the free-living, flavonoid-induced state ten years ago with non-strand-specific RNA-seq (Pérez-Montaño et al., 2016a, 2016b). Because the present experiment re-examines the same two conditions with strand-specific libraries and the same free-living reference, the two generations of data can be compared directly, and the comparison sets the interpretive baseline for everything that follows. Comparing them gene by gene first requires reconciling the gene identifiers, which have changed in both strains since 2016: in HH103 because the genome has been closed and reannotated (HH103_v32, Fuentes-Romero et al., 2025), and in CIAT 899 because the same 2012 assembly has since been reannotated by RefSeq, which renamed every locus tag. Of its 6,382 genes, 6,140 can be traced back to the identifiers used in 2016 and 242 are new gene calls without a counterpart there.

The first difference between the two generations is sensitivity. Genistein elicited 106 differentially expressed genes in HH103 in 2016 and 489 here, a 4.6-fold increase; apigenin elicited 19 in CIAT 899 in 2016 and 274 here, a 14.4-fold increase. Effect sizes also grew: for the genes both studies detect, the median absolute log_2_ fold change rose from 2.56 to 4.61 in HH103 and from 2.58 to 3.58 in CIAT 899. The second difference is that this gain is not spread evenly. The 2016 studies had sorted their responsive genes by what lay upstream of them. A gene preceded by a *nod* box or a *tts* box has an identified reason to answer the flavonoid, because NodD or TtsI binds there; a gene with neither had no such explanation, and appeared in those datasets without any link to the signalling cascade. That distinction turns out to predict almost perfectly which genes reproduce. In HH103, 100 of the 101 catalogued ORFs could be located in the current annotation and 87 were recovered as differentially expressed, but the recovery rate differs sharply by class (Figure 1d): 35 of 35 genes downstream of a *tts* box and 33 of 35 downstream of a *nod* box, against only 19 of 30 of the genes with no assigned element (Table S10). CIAT 899 gives the same result without a grey zone (Figure 1e). Reapplying the criterion of the earlier study to its published whole-genome table returns the 19 apigenin-responsive genes it reported, 16 of which have a counterpart in the current annotation: 12 of the 12 lying in a *nod* box operon are recovered here, and none of the 4 outside one. Its 20 qRT-PCR-validated genes behave the same way: all 15 that had been called induced at least three-fold are recovered and keep their ranking by induction strength, whereas of the five weakly regulated genes only two are recovered and three change signs (Table S11).

**Figure 1.**
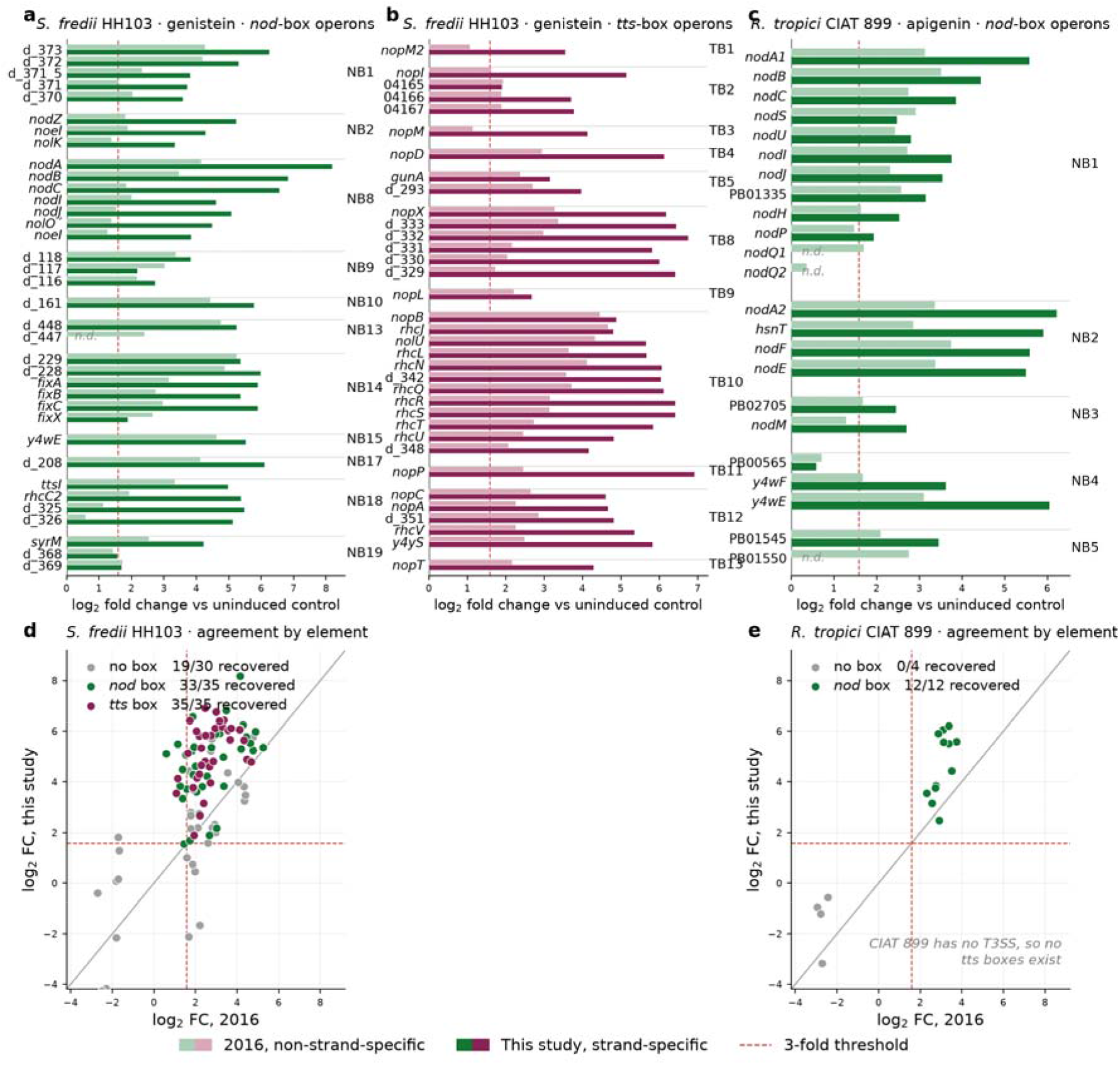
Fold changes of the genes of the *cis*-element-controlled operons, in the 2016 non-strand-specific experiments and in the present strand-specific data, grouped by regulatory element. Top row: (a) *nod* box and (b) *tts* box operons of HH103 under genistein, and (c) *nod* box operons of CIAT 899 under apigenin. The dashed line marks the three-fold threshold used to call differential expression. Genes without a counterpart in the current annotation are marked n.d. Values for 2016 are the inductions reported in the supplementary tables of Pérez-Montaño et al. (2016a, 2016b). Bottom row: agreement between the two generations, coloured by the regulatory element assigned in the 2016 study, for all 100 located ORFs of the HH103 catalogue (d) and the 16 located genes of the CIAT 899 catalogue (e); the grey line is identity and the legend gives the fraction of each class recovered as differentially expressed here. CIAT 899 encodes no type III secretion system, so it has no *tts* box counterpart to panel b. *Statistics*: this panel set is descriptive and carries no inferential test; the dashed line is the three-fold threshold (fold-change ≥ 3, log_2_ ≥ 1.585) and the recovery fractions in the legends of panels d and e are counts of genes reaching that threshold with an adjusted *P* < 0.005 in the present data. *Colour*: green marks genes under a *nod* box and wine red genes under a *tts* box, in each pair the lighter shade being the 2016 value and the darker one the present study; grey marks genes to which the 2016 studies assigned no regulatory element.

Seen operon by operon, the recovery of the *cis*-element-driven genes is complete (Figure 1a-c). Across the eleven *nod* box and eleven *tts* box operons of HH103 and the five active *nod* box operons of CIAT 899, not one operon is missing, and what the new data add is a uniform upward shift rather than a rearrangement: 84 of the 90 genes for which both studies report a value show a larger change now, and none changes sign. The median log_2_ fold change rises from 2.54 to 5.12 in the HH103 *nod* box genes, from 2.49 to 5.35 in the *tts* box genes and from 2.65 to 3.58 in the CIAT 899 *nod* box genes, and the number of these genes clearing the three-fold threshold rises from 78 to 88. The shift is largest where the 2016 values sat just above the detection limit, *nodA* of HH103 moves from 4.1 to 8.2 in log_2_ units and *nopP* from 2.5 to 6.9, while genes already strongly induced move comparatively little. Operons that the earlier data resolved only in part now appear induced along their full length: in *tts* box 10, the twelve genes spanning *nopB*, the *rhc* apparatus and *nolU* all exceed five-fold, and in *nod* box 8 the whole *nodABCIJ*-*nolO*-*noeI* stretch does the same.

Plotting each gene’s 2016 value against its value here makes the contrast between the two classes explicit (Figure 1d, e). The genes with an assigned *cis* element form one tight group lying above the line of equality, which means the older protocol had them right in direction and merely underestimated how strongly they respond. The 30 HH103 genes with no assigned element behave quite differently: they fall on both sides of the line, only 14 lie above it, six are now regulated in the opposite direction to what was reported, and seven no longer pass the three-fold threshold at all. In CIAT 899 the separation is absolute, the *nod* box genes occupying the upper right quadrant and the four genes without a box the lower left, none of them differentially expressed in the new data. These discrepancies do not reflect minor quantitative shifts, but rather unsubstantiated individual calls confined exclusively to the single class of genes for which no regulatory mechanism has ever been proposed.

Thus, the genes under an identifiable *cis* element, responding strongly, are insensitive to the library chemistry and reproduces in full, with its amplitude revised upwards. The weakly regulated genes that the earlier studies could not attribute to any regulator are where the two technologies disagree, and the most likely reason is the one those studies could not control for: without strand information, reads from a convergent neighbour or an antisense transcript are assigned to the wrong gene, which generates apparent regulation of modest size and arbitrary direction. Much of that unassigned class was therefore probably never part of the flavonoid response.

### 3.2 The free-living flavonoid response is confined to symbiotic functions

Before turning to the bacteroid, it is worth establishing what the flavonoid response looks like when resolved by function, since it is the baseline against which the symbiotic stage is read. Two complementary classifications are used throughout. The first, referred to here as functional blocks, is a purpose-built scheme that places every gene of each genome in exactly one of sixteen categories defined in Table 1; it covers 100% of the coding genes and is designed to make the symbiosis-specific machinery visible as such, at the cost of being specific to this pair of organisms. The second is the Clusters of Orthologous Groups classification, a standard, organism-independent assignment of proteins to broad functional categories by orthology, which covers 76% of the genes here and allows the results to be read against the wider bacterial literature, at the cost of dissolving the symbiotic genes into generic categories or leaving them unassigned. Both were tested the same way, by a two-sided Fisher exact test of each category against the rest of the genome with Benjamini-Hochberg correction within each condition (Tables S4 and S8). Over- and under-representation are always relative to the response as a whole: a category is over-represented when it contributes more differentially expressed genes than its share of the genome would give if those genes were drawn at random, and under-represented when it contributes fewer. The blocks themselves, with the content of each and its size in the two genomes, are given in Table 1.

The block analysis shows that in both strains the response is narrow and concentrated on the machinery of recognition, but that the two strains differ in what they switch off (Figure 2).

**Figure 2.**
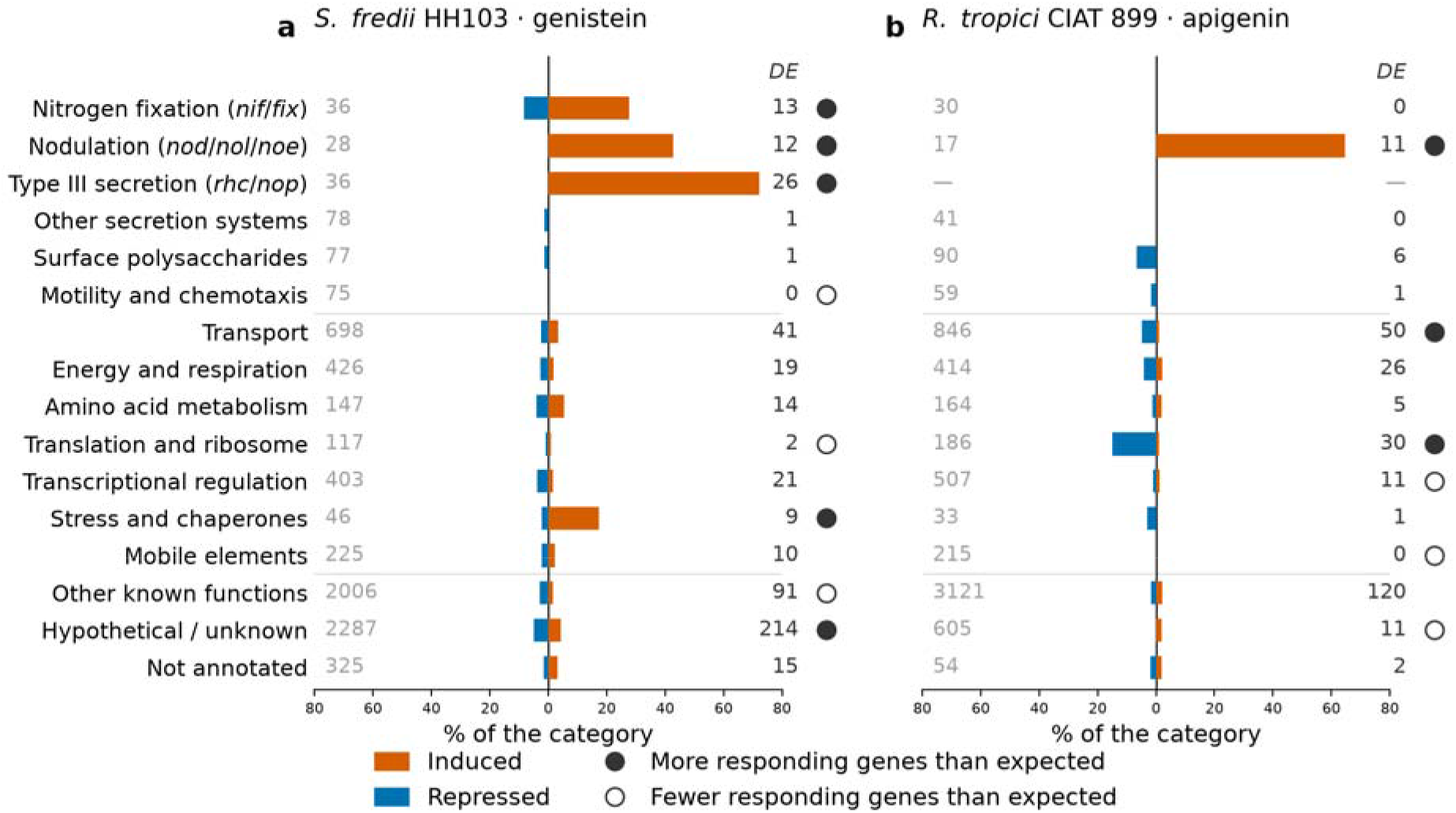
Functional blocks of the free-living flavonoid response. (a) *S. fredii* HH103 induced with genistein; (b) *R. tropici* CIAT 899 induced with apigenin. Bars give the percentage of the genes of each block that are differentially expressed, induced to the right in orange and repressed to the left in blue; the grey number at the left of each row is the number of genes the block contains in that genome and the number at the right, is how many of them respond (DEGs, fold-change ≥ 3, log_2_ ≥ 1.585). Circles mark blocks whose number of responding genes departs from expectation: filled, more than expected; open, fewer than expected (The expectation is what the block would contain if the differentially expressed genes were a random sample of the genome). These circles describe how many genes respond, not the direction in which they change. *Statistics*: for each block, a two-sided Fisher exact test of responding against non-responding genes in that block versus the rest of the genome, with the Benjamini-Hochberg correction applied across the sixteen blocks of HH103 and the fifteen of CIAT 899 within each condition; circles are drawn at FDR < 0.05. The universe is 7010 genes for HH103 and 6382 for CIAT 899. Blocks absent from a genome are marked with a dash: CIAT 899 encodes no type III secretion system. Full statistics in Table S4.

In HH103, genistein mobilises 26 of the 36 *rhc* and *nop* genes of the type III secretion system (72% of the category, Odds Ratio (OR) 36.6) and 12 of the 28 nodulation genes (43%, OR 10.2), and in both categories every single responding gene is induced, without one exception. Nitrogen fixation follows at 13 of 36 genes (36%, OR 7.7; 10 induced, 3 repressed) and the stress and chaperone block at 9 of 46 (20%, OR 3.3; 8 induced, 1 repressed). The complement is as informative: not one of the 75 flagellar and chemotaxis genes responds at all, translation is under-represented (2 of 117; OR 0.23), and the other secretion systems and the surface polysaccharide genes contribute one repressed gene each. The flavonoid signal, in other words, reaches the two secreted signalling systems and almost nothing else.

CIAT 899 shows the same concentration on nodulation, 11 of its 17 *nod*, *nol* and *noe* genes (65%, OR 42.5), again all induced. Lacking a type III secretion system, it has no second induced block to deploy; instead apigenin represses. Translation and ribosome biogenesis is over-represented (30 of 186 genes, OR 4.7) and 28 of those 30 are repressed; transport is over-represented (50 of 846, OR 1.5) with 42 of 50 repressed; and the surface polysaccharide genes that respond are repressed without exception. Mobile elements are excluded altogether, none of the 215 transposases and integrases responding. Overall, 51% of the HH103 genistein response is inductive against 43% of the CIAT 899 apigenin response.

Set side by side, the two responses share a narrow common ground and diverge everywhere else. Nodulation is the only category over-represented in both strains, and in both it responds by induction alone. The categories each strain leaves alone are also the same: the flagellar and chemotaxis genes are essentially untouched by either flavonoid (0 of 75 in HH103, 1 of 59 in CIAT 899), and the surface polysaccharide genes contribute only repressed genes in both (1 and 6). Beyond that the strategies have little in common. HH103 adds to the induced side, pre-activating the fixation machinery and the chaperone block alongside the type III system. CIAT 899 does neither, not one of its 30 *nif*/*fix* genes responds to apigenin, and its stress block contributes a single repressed gene, and instead builds its response on the repressed side: translation (28 of the 30 responding genes repressed), transport (42 of 50) and the complete exclusion of mobile elements, with none of its 215 transposases and integrases responding. The genomic location of the two regulons tells the same story. In HH103, 191 of the 489 genistein-responsive genes (39.1%) lie on the symbiotic plasmid pSfHH103d, against 180 on the chromosome and 100 on the megaplasmid pSfHH103e, so the symbiotic replicon carries the largest share of the response despite holding less than a tenth of the genome. CIAT 899 does not concentrate its response in the same way: only 26 of the 274 apigenin-responsive genes (9.5%) lie on its symbiotic plasmid pRtrCIAT899b, which carries 8.1% of the genes in the genome, so the symbiotic replicon contributes no more than its share. The *nod* genes it induces are indeed there, all five *nodD* and all three *nodA* copies are on pRtrCIAT899b, but they are few enough that the replicon as a whole shows no excess, and the bulk of the apigenin response lies on the chromosome (148 of 274, 54.0%).

The COG classification adds little to the HH103 flavonoid response and a great deal to the CIAT 899 one, for a reason that is itself informative. Of the 489 genistein-responsive genes, 326 carry a COG assignment, yet only one category is over-represented among them, inorganic ion transport and metabolism (P: 36 of 315 genes, OR 2.11, FDR 0.005). The type III secretion and nodulation genes that dominate the block analysis are simply invisible at this resolution, because *rhc*, *nop* and *nod* genes either fall into broad categories or carry no COG at all, the same limitation that hides the *nif*/*fix* signal. In CIAT 899, by contrast, 222 of the 274 apigenin-responsive genes carry a COG and three categories are significant: translation, ribosomal structure and biogenesis is over-represented (J: 31 of 245 genes, OR 3.23), while signal transduction (T: OR 0.21) and transcription (K: OR 0.43) are under-represented. The repression of the translational machinery therefore reproduces on an orthology-based classification that knows nothing about symbiosis, which makes it a property of the cell’s physiology rather than an artefact of how the symbiotic categories were drawn.

The difference between the two strains is therefore not one of sensitivity to the flavonoid but of what each does with the signal. HH103 answers by deploying a secretion system and priming the machinery of fixation, with a large accompanying set of genes whose role remains unknown. CIAT 899 answers by inducing nodulation genes alone and damping its translational and transport capacity while it builds Nod factors, on a regulon that is small, almost entirely annotated and almost entirely accounted for by the *nod* box architecture.

### 3.3 A shared core dominates the bacteroid response, and host identity rather than nodule type shapes the rest

Strand-specific RNA-seq of bacteroids recovered from three legume hosts of *S. fredii* HH103 (*G. max*, *G. uralensis* and *L. burttii*) and two hosts of *R. tropici* CIAT 899 (*P. vulgaris* and *L. burttii*), each referred to the free-living control of its own strain, identified between 1,877 and 2,284 DEGs per host-strain combination (fold change of at least 3, log2 ≥ 1.585, adjusted P < 0.005; Table 2, Table S1). Across both strains the bacteroid state therefore mobilises roughly a third of the protein-coding complement. The two strains differ consistently in the balance of the response: every HH103 condition is induction-leaning, at 53.9 to 55.7% of its DEGs, and every CIAT 899 condition sits at 43.3 to 43.7% (Table 2).

**Table 2.** Conditions analysed and size of each response. DEGs were called with DESeq2 against free-living control cells of the same strain, by the Wald test with Benjamini-Hochberg correction, and retained at a fold change of at least 3 (log_2_ ≥ 1.585) and an adjusted *P*<0.005; three libraries per condition. Host-exclusive DEGs are those responding in that host and in no other host of the same strain. *L. burttii* is nodulated by both strains and is the only host shared between them.

| **Strain** | **Condition** | **Nodule type** | **Relationship to strain** | **DEGs** | **Induced** | **Repressed** | **Induced (%)** | **Host-exclusive** |
|---|---|---|---|---|---|---|---|---|
| **HH103** | Free-living + genistein | - | - | 489 | 250 | 239 | 51.1 | - |
| **HH103** | *Glycine max* (soybean) | Determinate | Natural host | 1,914 | 1,055 | 859 | 55.1 | 435 |
| **HH103** | *Lotus burttii* | Determinate | Extended range | 2,104 | 1,171 | 933 | 55.7 | 337 |
| **HH103** | *Glycyrrhiza uralensis* | Indeterminate | Extended range | 1,877 | 1,012 | 865 | 53.9 | 214 |
| **CIAT 899** | Free-living + apigenin | - | - | 274 | 119 | 155 | 43.4 | - |
| **CIAT 899** | *Phaseolus vulgaris* (common bean) | Determinate | Natural host | 2,083 | 902 | 1,181 | 43.3 | 419 |
| **CIAT 899** | *Lotus burttii* | Determinate | Extended range | 2,284 | 997 | 1,287 | 43.7 | 620 |

Partitioning these genes by host shows that the shared component is by far the largest (Figure 3; Tables S2 and S3). In HH103, 1,171 genes were differentially expressed in all three hosts, against 435, 337 and 214 exclusive to *G. max*, *L. burttii* and *G. uralensis*, respectively. In CIAT 899, 1,664 genes were shared between the two hosts, with 620 and 419 exclusive to *L. burttii* and *P. vulgaris*. The bacteroid programme is thus largely the same whichever legume the bacterium ends up in, and only a minority of it is host-restricted. In HH103 that conclusion rests on a deliberately demanding criterion: requiring a gene to respond in all three hosts leaves 1,171 of the 2,855 genes of the union, 41.0%, whereas the same calculation over soybean and *L. burttii* alone would return 1,377 of 2641, 52.1%. *G. uralensis* forms indeterminate nodules and *G. max* and *L. burttii* determinate ones, so if nodule ontogeny were the organising axis the two determinate hosts should resemble each other and *G. uralensis* stand apart. The partitions say the opposite (Figure 3a): *G. uralensis* and *L. burttii* share 390 DEGs to the exclusion of soybean, whereas *G. max* shares only 206 with *L. burttii* and 102 with *G. uralensis*; and the largest host-exclusive set belongs to *G. max* (435 genes) while the smallest belongs to the indeterminate-nodule host *G. uralensis* (214 genes). Soybean, the natural and coevolved host of HH103, is the outlier among the three, and the two legumes of its extended range behave alike. In *S. fredii* HH103, what differentiates the rhizobial side of the interaction is therefore the evolutionary relationship with the partner, not the developmental programme of the organ.

**Figure 3.**
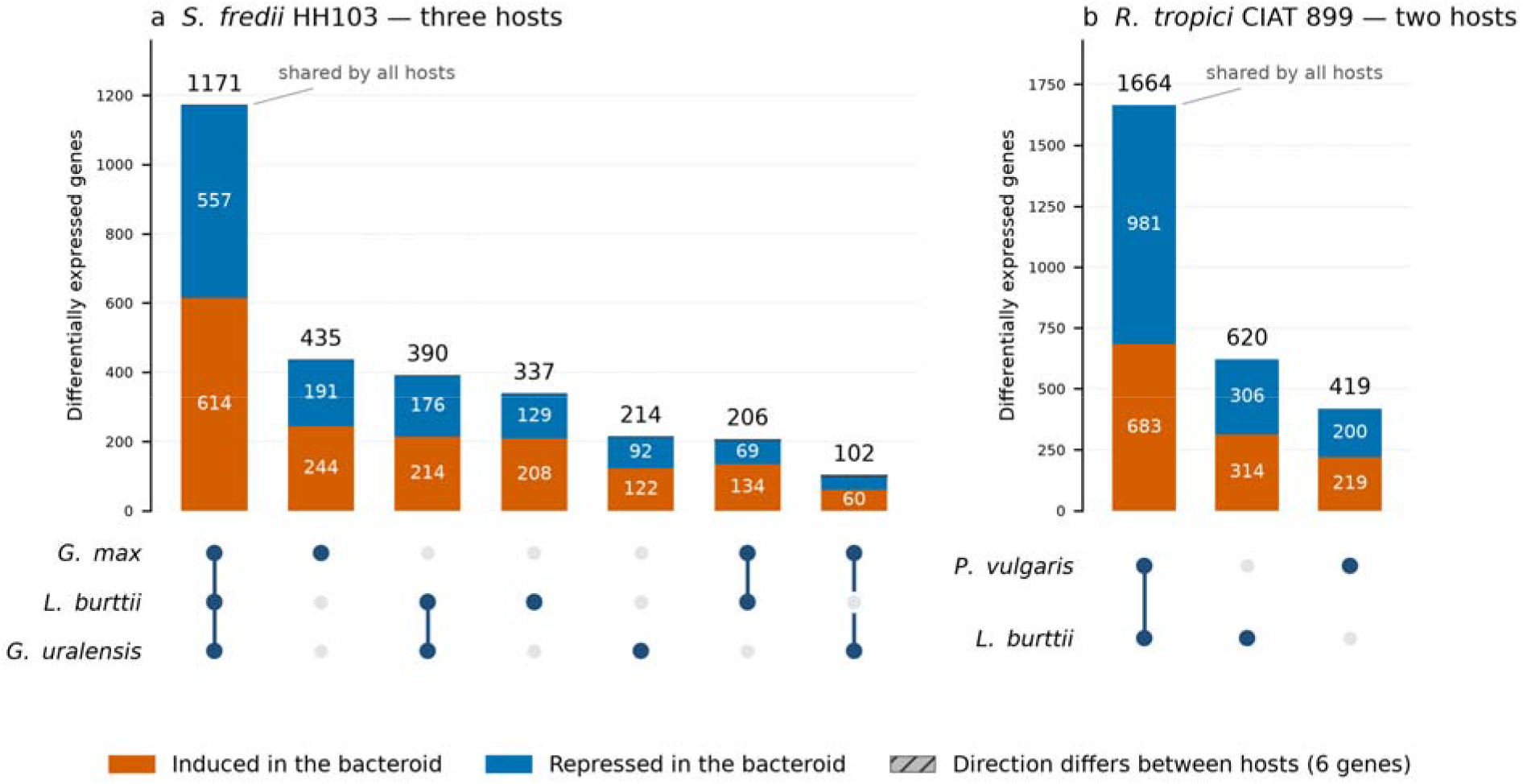
Host-specificity partitions of differentially expressed genes, with each partition split by the direction of the change. (a) *S. fredii* HH103 across three hosts; (b) *R. tropici* CIAT 899 across two hosts. The matrix below each panel marks which hosts make up each partition; hosts forming determinate nodules are listed first (*G. max* and *L. burttii*; *P. vulgaris* and *L. burttii*) and the indeterminate-nodule host *G. uralensis* last, and a connector crossing an open circle skips that host. Bars are stacked as induced over repressed, with the partition total above each bar. Six HH103 genes are regulated in opposite directions in different hosts and form the hatched segments of the *G. max*+*L. burttii* and *G. max*+*G. uralensis* partitions (three each). *Statistics*: partition membership is a set operation on the DEG lists, with no inferential test; a gene enters a partition when it passes the fold change ≥ 3 (log_2_ ≥ 1.585) and adjusted *P* < 0.005 criterion in exactly the hosts marked. *Colour*: orange is induced and blue repressed in the bacteroid, as throughout.

Every HH103 partition leans towards induction, least in the shared core (614 of 1,171 genes induced, 52.4%) and most in the smaller host-restricted sets (*G. max*+*L. burttii*, 66.0%; *L. burttii*-exclusive, 61.7%; *G. max*+*G. uralensis*, 60.6%), with the remaining sets between 54.9% and 57.0%. CIAT 899 inverts the pattern where it matters most: its shared core is the only partition in either strain in which repression predominates, 981 of 1,664 genes down against 683 up (41.0% induced), while its two host-exclusive sets sit at 50.6% and 52.3%, as close to even as any partition here. Repression is therefore not a by-product confined to the host-specific fraction but an integral part of the common bacteroid programme, and in CIAT 899 the larger half of it. Moreover, host identity acts almost entirely on whether a gene is mobilised rather than on the direction of its change. Of the 2,855 HH103 genes differentially expressed in at least one host, only 6 changed signs between hosts; among the 2,703 CIAT 899 genes, none did.

### 3.4 Functional blocks in the bacteroid: fixation and stress up, motility down, type III secretion host-dependent

Applying the same test to the five bacteroid conditions gives a picture that is consistent across hosts and, in its main features, across strains (Figure 4, Table S4). Two categories are over-represented in all five: nitrogen fixation, with 58-87% of the *nif*/*fix* genes differentially expressed and essentially all of them induced (OR 3.3-13.6), and the stress and chaperone block, at 43-64% of its genes and OR 2.1-3.2. Transport is over-represented in four of the five conditions.

**Figure 4.**
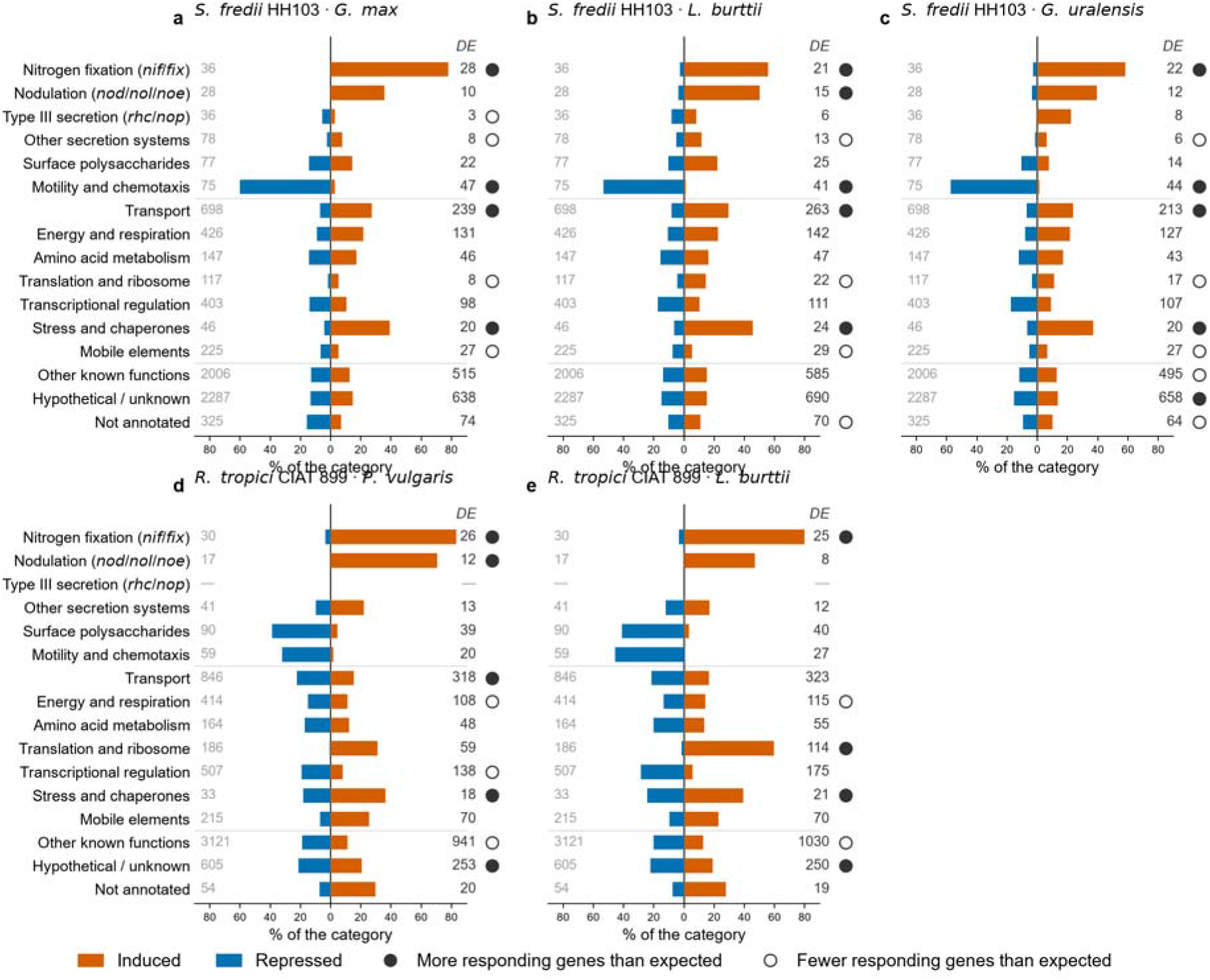
Functional blocks of the bacteroid response. (a-c) *S. fredii* HH103 bacteroids from *G. max*, *L. burttii* and *G. uralensis*; (d, e) *R. tropici* CIAT 899 bacteroids from *P. vulgaris* and *L. burttii*. Bars give the percentage of the genes of each block that are differentially expressed, induced to the right in orange and repressed to the left in blue; the grey number at the left of each row is the number of genes the block contains in that genome and the number at the right, is how many of them respond (DEGs, fold-change ≥ 3, log_2_ ≥ 1.585). Circles mark blocks whose number of responding genes departs from expectation: filled, more than expected; open, fewer than expected (The expectation is what the block would contain if the differentially expressed genes were a random sample of the genome). These circles describe how many genes respond, not the direction in which they change. *Statistics*: two-sided Fisher exact test per block against the rest of the genome, Benjamini-Hochberg correction across the blocks within each condition, circles at FDR < 0.05; universe 7010 genes for HH103 and 6382 for CIAT 899. Full statistics in Table S4.

The categories that move downwards are the ones that matter for the comparison with the free-living stage. Motility and chemotaxis is the most strongly repressed block in every bacteroid: between 34% and 63% of the 75 HH103 and 59 CIAT 899 flagellar and chemotaxis genes are differentially expressed, and the repressed fraction accounts for 45 of 47 in *G. max*, 43 of 44 in *G. uralensis*, 40 of 41 in *L. burttii*, and all 27 in CIAT 899 bacteroids from *L. burttii*. The other secretion systems, type I, type II, type IV and the Sec and Tat translocases, are under-represented in all three HH103 bacteroids (OR 0.23-0.46), and mobile elements are depleted in the same three (OR ∼0.35). The contrast with the free-living response is direct: genistein switches the type III system on and leaves motility untouched, whereas inside the nodule the flagellar genes are shut down and most of the secretion machinery falls silent.

Interestingly, the T3SS itself is the exception, since it behaves differently in the indeterminate-nodule host. As a category it is never over-represented in a bacteroid, and in *G. max* it is clearly under-represented (3 of 36 genes, OR 0.24); but the direction of the few genes that do respond splits the hosts cleanly. In *G. max* only three *rhc*/*nop* genes respond and two of them are repressed, *nopA* (log_2_ -2.40) and *nopC* (-2.64), with *nopP* the only induced one; in *L. burttii* six respond and the same three structural genes are repressed, *nopB*, *nopA* and *nopC*, against *nopP* and *rhcC2* induced. In *G. uralensis* eight genes of the block respond and every one of them is induced: *rhcC2*, both copies of *nopP*, and the effectors *nopM*, *nopX*, *nopT* and *nopD*, together with one conserved hypothetical protein assigned to the block. The distinction is also structural: what the determinate hosts repress are the surface and translocon components NopA, NopB and NopC, whereas what the indeterminate host induces are the translocated effectors. Nops are known to act as symbiotic factors on *G. uralensis* and on *G. max* but as asymbiotic ones on other legumes (López-Baena et al., 2008; Jiménez-Guerrero et al., 2022), and these data indicate that the difference extends into the nitrogen-fixing stage: in the indeterminate nodules of *G. uralensis* the effector repertoire is still being expressed when fixation is under way, while in the determinate nodules of soybean and *L. burttii* the apparatus that delivers it is being dismantled.

The two strains also differ in which replicon supplies the bacteroid programme, and they differ in opposite senses. In HH103 the symbiotic plasmid pSfHH103d, which holds 9.5% of the genes and carried 39.1% of the genistein response, falls to 11.2-13.4% of the bacteroid DEGs (236 of 2,104 in *L. burttii*, 246 of 1,914 in *G. max*, 252 of 1,877 in *G. uralensis*), while the chromosome and pSfHH103e take over. In CIAT 899 the symbiotic plasmid pRtrCIAT899b moves the other way: from 9.5% of the apigenin response, which is its share of the genome, to 15.7% of the bacteroid DEGs in *P. vulgaris* (327 of 2083) and 12.2% in *L. burttii* (278 of 2,284). The symbiotic replicon is thus over-represented in the free-living flavonoid response of HH103 and in the bacteroid response of CIAT 899, the reverse of each other, which is consistent with what each replicon carries: a flavonoid-inducible type III secretion system in HH103, and in CIAT 899 a nod gene complement whose products are needed throughout the interaction.

One block diverges between the strains. Translation and ribosome biogenesis is under-represented in all three HH103 bacteroids (8 to 22 of 117 genes; OR 0.19-0.54) but strongly over-represented in CIAT 899 bacteroids from *L. burttii*, where 114 of 186 genes respond and 111 of them are induced (OR 2.9); in *P. vulgaris* bacteroids the same block is mobilised at 32% with 58 of 59 genes induced although the enrichment itself is not significant. The two strains therefore arrive at the nitrogen-fixing state with opposite settings for their protein-synthesis capacity.

An independent enrichment over COG functional categories reproduces these conclusions and adds further detail (Tables S8 and S9). Cell motility (category N) is the most strongly over-represented category of the HH103 bacteroids, 46, 42 and 40 of its 74 genes in *G. max*, *G. uralensis* and *L. burttii* (OR 4.5, 3.7 and 2.7), with 42, 38 and 36 of those genes repressed: the COG counterpart of the flagellar silencing described above. Translation (J) is under-represented in the same three bacteroids (OR 0.44–0.62), where the few genes that respond divide evenly between induction and repression, and also in *P. vulgaris* (OR 0.57), where 41 of the 52 responding genes are induced; in *L. burttii* bacteroids of CIAT 899 it is over-represented instead, at 110 of 245 genes (OR 1.54) with 94 of them induced. The two CIAT 899 hosts therefore differ in how much of the translational machinery they mobilise rather than in the direction it takes, and both differ from HH103. Replication, recombination and repair (L) is under-represented in the HH103 bacteroids (OR 0.58–0.64) and the genes that do respond are mostly repressed (33 of 41, 28 of 42 and 29 of 45), consistent with the repression of mobile elements. Two signals appear only in the COG analysis. Carbohydrate transport and metabolism (G) is over-represented in both CIAT 899 bacteroids, 211 of 480 genes in *P. vulgaris* and 227 in *L. burttii* (OR 1.82 and 1.76), and predominantly repressed in both (160 and 163 of the responding genes), whereas in HH103 it responds at the background rate (OR 1.03–1.25) and mostly upwards. Extracellular structures (W) is over-represented in both CIAT 899 bacteroids (12 and 13 of 19 genes; OR 3.7 and 4.0) but, among the HH103 bacteroids, only in *L. burttii* (23 of 46; OR 2.3), the enrichment falling short of significance in *G. uralensis* (OR 2.1) and absent in *G. max* (OR 1.05).

### 3.5 Most of the recognition programme survives into the nodule, but the bacteroid programme is largely new

Because the bacteroid and flavonoid samples share the same free-living reference, the whole trajectory can be followed gene by gene within each strain, and because 3,347 orthologous pairs could be defined between the two proteomes by reciprocal best hits (median amino-acid identity 71.5%; Table S5), the two trajectories can also be compared, at least in some extend, with each other.

The first result is how much of the recognition programme survives into the nodule. In HH103, 378 of the 489 genistein-responsive genes (77.3%) are also differentially expressed in at least one bacteroid, 231 of them in the three-host core, leaving only 111 exclusive to the free-living condition; and where the two overlap they agree on direction, in 94.4% of cases in *G. uralensis* (303/321), 92.4% in *L. burttii* (291/315) and 87.9% in *G. max* (247/281). CIAT 899 carries less forward and less faithfully: 168 of its 274 apigenin-responsive genes (61.3%) are also bacteroid DEGs, with 83.7% directional concordance in *P. vulgaris* (113/135) but only 65.6% in *L. burttii* (84/128). Part of the early response is therefore not switched off on entry but sustained into the fixing nodule, and more so in HH103 than in CIAT 899. The persistence is clearest for the canonical regulon itself: of the 35 top genistein-responsive genes of Pérez-Montaño et al. (2016a), 18 remain differentially expressed in *G. max* bacteroids, 23 in *G. uralensis* and 18 in *L. burttii*, almost all of them still induced (Table S6). The recognition machinery is not dismantled once the bacterium is inside.

Following each flavonoid-induced operon into the nodule shows that the dismantling is selective rather than general, and that what survives depends on both the operon and the host (Figure 5). Three patterns separate cleanly. The type III secretion apparatus, but not the related-effectors, is switched off outright: of the twelve genes of *tts* box 10, which span *nopB*, *nolU* and the *rhcJLNQRSTU* core, not one remains differentially expressed in *G. max* or *G. uralensis* bacteroids and only one does in *L. burttii*, despite all twelve being induced 18-to 85-fold by genistein. The Nod factor operons are attenuated but retained: *nod* box 8, carrying *nodABCIJ*-*nolO*-*noeI*, keeps five to seven of its seven genes differentially expressed in every host, with a median drop of 2.1 to 3.7 log_2_ units, and in CIAT 899 the two Nod factor operons behave the same way, *nod* box 1 keeping 7 of 10 genes in *P. vulgaris* and *nod* box 2 all four, both attenuated by 1.5 to 3.2 units. And one operon is not attenuated at all but amplified: *nod* box 14, which carries *fixABCX* together with an iron-sulfur cluster assembly protein, keeps all six genes in all three hosts and rises by a median of 2.6 to 4.2 log_2_ units, *fixA* and *fixC* reaching 10.4 and 10.1 in soybean bacteroids against 5.9 in free-living cells. A *nod* box therefore does not by itself predict that a gene belongs to the early programme: when it controls fixation genes, the operon behaves like the bacteroid programme.

**Figure 5.**
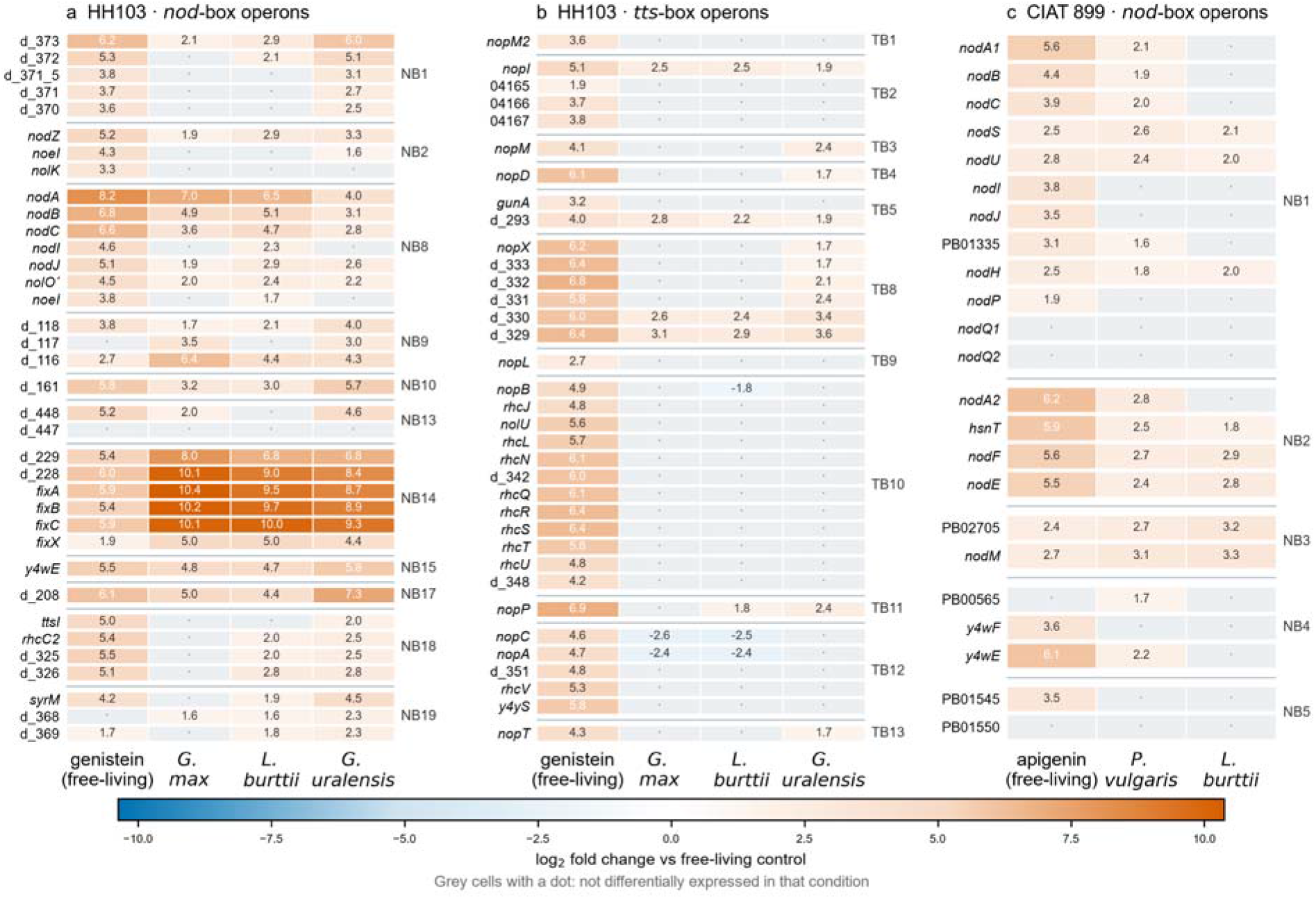
Fate of the flavonoid-induced operons inside the nodule. Each cell is the log_2_ fold change of one gene against free-living control cells, for the free-living flavonoid condition and for each bacteroid host; grey cells with a dot mark genes not differentially expressed in that condition. (a) *nod* box and (b) *tts* box operons of HH103; (c) *nod* box operons of CIAT 899. Rows are grouped by regulatory element, hosts forming determinate nodules precede the indeterminate-nodule host *G. uralensis*, and gene names are given where assigned and locus tags otherwise. *Statistics*: each value is the DESeq2 log_2_ fold change for that contrast; a cell is drawn in colour when the gene passes fold change ≥ 3 (log_2_ ≥ 1.585) and adjusted *P* < 0.005 in that condition and grey with a dot when it does not. The colour scale is continuous and centred on zero, orange for induction and blue for repression, as throughout; no test is applied to the operons as groups.

The host dependence runs along the same axis identified for the Nops. In *G. uralensis*, *nod* box 1 retains all five of its genes and *nod* box 18 all four, against one of five and none of four in soybean; and the effector-bearing *tts* boxes behave likewise, with *tts* box 8 keeping all six genes in *G. uralensis* but only two in the two determinate hosts, and *tts* boxes 3, 4 and 13 retaining their gene in *G. uralensis* alone. Conversely *tts* box 12, which carries *nopC* and *nopA*, is the one place where the determinate hosts do something the indeterminate host does not: both genes are repressed in *G. max* and *L. burttii* by about seven log_2_ units relative to the genistein condition, and neither responds in *G. uralensis*. In CIAT 899 the equivalent asymmetry favours the natural host, *nod* box 1 retaining 7 of 10 genes in *P. vulgaris* against 3 of 10 in *L. burttii*.

Seen from the other end, however, the bacteroid programme is overwhelmingly new. Of the 2,966 HH103 and 2,809 CIAT 899 genes responding in at least one condition, 2,477 and 2,535 respectively are exclusive to the bacteroid state; only 378 and 168 respond in both states, and 111 and 106 respond to the flavonoid alone (Figure 6a). The transition is thus not a continuation of the early dialogue but the superimposition of a far larger programme on a residue of it.

**Figure 6.**
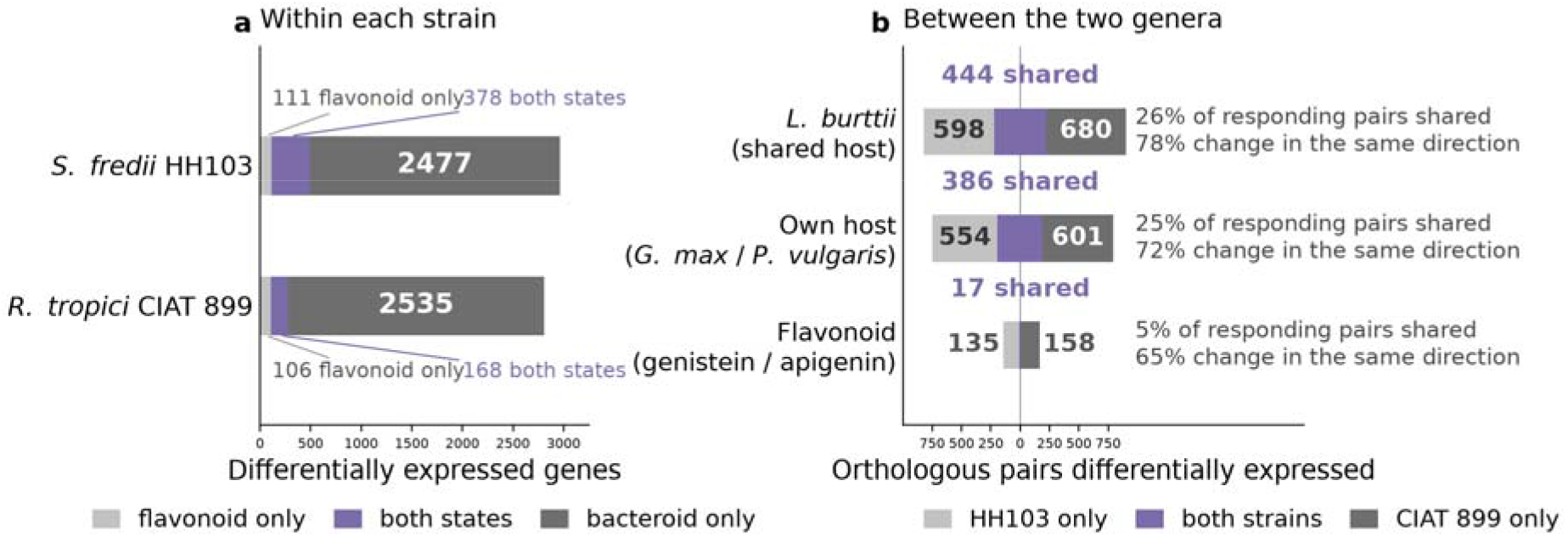
Recognition is strain-private, differentiation is shared. (a) Within each strain, overlap between the genes responding to the flavonoid in free-living cells and those responding in bacteroids (union of all hosts of that strain). (b) Between the two strains, fate of the 3347 reciprocal best-hit orthologue pairs: for each comparison the bar shows how many pairs are differentially expressed in HH103 only (blue, extending left), in CIAT 899 only (orange, extending right) and in both strains (purple, straddling the centre). Comparisons are the shared host *L. burttii*, each strain’s own host, and the free-living flavonoid condition; beside each bar are the percentage of responding pairs shared by the two strains and, of those, the percentage changing in the same direction. *Statistics*: all quantities are counts and percentages over the 3347 reciprocal best-hit orthologue pairs (BLASTP, at least 30% amino-acid identity over at least 50% of the query), with no inferential test. *Colour*: purple marks the genes or pairs responding in both states or in both strains, and the two greys the sets private to one state or one strain; this figure carries no direction information, so the orange and blue of the other figures do not appear in it.

Comparing the two strains to each other requires a common currency, and the 3,347 reciprocal best-hit orthologue pairs between the two genomes provide it: each pair is one gene of HH103 together with its closest counterpart in CIAT 899, so a pair can be differentially expressed in HH103 only, in CIAT 899 only, or in both. Asking how often the two bacteria move the same pair gives a direct measure of how much of each programme is conserved between the two genera. In the shared host *L. burttii*, 1,722 pairs respond in at least one strain and 444 of them (26%) respond in both, 78% of those in the same direction. On each strain’s own host the figure is almost identical: of 1,541 responding pairs, 386 (25%) respond in both, 72% concordantly. The free-living flavonoid responses behave completely differently: only 310 pairs respond at all, and a mere 17 of them (5%) in both strains, with 65% directional agreement (Figure 6b). The bacteroid programme is therefore conserved across genera to a quarter of its extent, whereas the flavonoid-induced programme is almost entirely strain-private.

Taken together, the two ends of the trajectory describe the same transition. The flavonoid response is strain-private, built on a secretion system in one strain and on signal chemistry diversity in the other. The bacteroid response is largely shared, with the fixation and chaperone blocks engaged in every host and the flagellar genes silenced. Each bacterium therefore begins the symbiosis speaking its own language and ends it in a physiological state its partner would recognise, with host identity modulating the amplitude of that end state rather than its content.

## 4. Discussion

The central outcome of this work is the asymmetry between two stages of the symbiotic interaction. The flavonoid-induced programme, the chemical dialogue that precedes infection, is almost entirely private to each strain: of the 310 orthologue pairs that respond to a flavonoid in either bacterium, only 17 respond in both, and the two strains build that response out of different materials, a type III secretion system in one and a repression of translational and transport capacity in the other. The bacteroid programme, by contrast, converges on hundreds of orthologues across two genera and is dominated in both strains by nitrogen fixation and stress tolerance with motility shut down. Different rhizobia appear to negotiate entry in idiosyncratic dialects while converging on a largely shared physiological end state.

The genomic organisation of the two stages follows the same logic in each strain but in opposite directions, which is informative about what the symbiotic replicons are for. In HH103 the symbiotic plasmid carries 39.1% of the flavonoid response against the 9.5% of the genome it represents, and falls back to roughly its share in the bacteroid; in CIAT 899 the symbiotic plasmid is not over-represented in the apigenin response at all and becomes over-represented in the bacteroid. The difference tracks what each replicon encodes: a flavonoid-inducible secretion system whose job is done early, against a *nod* gene complement under five regulators whose products are needed throughout. What the HH103 symbiotic plasmid contributes to the bacteroid, moreover, is disproportionately the host-independent part of the programme, being enriched in the shared core and depleted from the host-exclusive sets.

A second difference in regulatory wiring concerns the fixation genes themselves, and it bears on the same point. In CIAT 899 not one of the 30 *nif*/*fix* genes lies in a *nod* box operon, and under apigenin the block is not merely below threshold but flat: the median log_2_ ratio across the 30 genes is -0.28 and the entire range falls between -0.87 and +1.58. A single chromosomal gene of unknown product passes the significance cut and falls just short of the three-fold threshold, so the block contributes no differentially expressed gene at all. In the nodule the same block is almost fully engaged, 26 of 30 genes in *P. vulgaris* and 25 in *L. burttii*, with median inductions of 8.4 and 6.7 log_2_ units. HH103 behaves differently at the early stage: 13 of its 36 *nif*/*fix* genes already respond to genistein, and five of them are the *fixABCX* block of *nod* box 14, with *fixA* and *fixC* already close to 6 log_2_ units in free-living cells and above 10 in soybean bacteroids. Since the end states are comparable, median inductions of 8.0 to 9.0 across the three HH103 hosts against 6.7 to 8.4 in the two CIAT 899 hosts, the strains differ in when the fixation machinery is first engaged rather than in the level it ultimately reaches.

The regulatory logic makes the contrast interpretable. In the rhizobia where this has been dissected, *fixABCX* and *nifHDK* are NifA-dependent and respond to low oxygen tension through the FixLJ two-component system, with *nifA* itself frequently autoregulated by read-through from the upstream *fixABCX* operon (Martínez et al., 2004). Placing *fixABCX* under a *nod* box, as HH103 does, adds a flavonoid- and NodD-dependent input to a circuit that is otherwise oxygen-controlled, which amounts to priming the electron supply to nitrogenase during the chemical dialogue, before any microoxic niche exists. That this is a strain-level acquisition rather than a general property of rhizobia is clear from CIAT 899, where the genomic opportunity exists but is not used: the *nod* box 5 operon and *nodD4* lie immediately upstream of *nifH* on the symbiotic plasmid, *nodD4* ending 606 bp before the *nifH* start, and yet the nitrogenase operon shows no flavonoid response whatever. A *nod* box is therefore not a marker of the early programme in itself, and in HH103 it is the route by which part of the late programme is brought forward.

The research was also designed to test whether determinate and indeterminate nodules impose distinguishable transcriptional programmes on the same rhizobial genome. Within HH103 they do not: the indeterminate-nodule host *G. uralensis* resembles *L. burttii* more closely than either resembles soybean. This extends to a three-host transcriptomic design the caution already raised on morphological grounds by Oono et al. (2010), and it rules out the most obvious alternative explanation for a null result of this kind. Where nodule type does reorganise bacteroid physiology it usually does so through terminal differentiation, and the three HH103 hosts do not differ in that respect, since HH103 bacteroids are not terminally differentiated even in the indeterminate, IRLC nodules of *G. uralensis* (Crespo-Rivas et al., 2016). Nodule architecture is therefore varied here while differentiation fate is held constant, and the transcriptome follows neither architecture nor, by implication, the NCR-driven route. The design further separates two variables that a two-host comparison necessarily confounds: with only one determinate and one indeterminate host, any difference between them can be read as an effect of nodule type, whereas with two determinate hosts it becomes clear that the larger split lies elsewhere. What the present data cannot say is whether a rhizobium whose bacteroids do differentiate terminally would behave the same way; that question needs a strain-host pair in which terminal differentiation is documented, and is the natural extension of this work.

Although nodule type does not organise the response as a whole, one block does distinguish the indeterminate host, and it is the one the literature would predict. The T3SS genes that respond in *G. uralensis* bacteroids are uniformly induced and all of them translocated effectors, whereas the genes that move in the two determinate hosts are the components of the surface appendage and the translocon, and they go down. Together with the persistence of 18 to 23 of the 35 canonical genistein-induced genes inside nodules, this argues against a clean switch from a nodulation programme to a fixation programme: the *nod* and type III blocks are attenuated rather than silenced, and *G. uralensis* retains the most. The host dependence of Nop action established for the infection stage therefore appears to extend into the nitrogen-fixing one, with effector delivery still apparently required during nodule maturation in at least one host.

Alongside nitrogen fixation, the chaperone and stress-response block is over-represented in every bacteroid condition examined, in both strains and on every host. The microaerobic, osmotically demanding environment of the symbiosome, and the proteostatic load of assembling nitrogenase, offer a straightforward explanation. The divergent behaviour of the translational machinery between the two strains, repressed in HH103, induced in CIAT 899 bacteroids from *L. burttii*, suggests that the two organisms reach the differentiated state by different routes, and deserves direct testing. Mobile elements move the other way: transposases and integrases are depleted among the DEGs of all three HH103 bacteroids and completely absent from the apigenin response of CIAT 899. Given the unusually high transposable-element content of the HH103 genome documented by Fuentes-Romero et al. (2025), the repression of this compartment inside the host is notable and may reflect selection against genome instability during the symbiotic phase.

In conclusion, following the same two genomes from the flavonoid-induced free-living cell to the mature bacteroid shows that the symbiosis is run as two programmes rather than one continuum. The early programme is strain-private and built from different materials in each bacterium; the bacteroid programme is large, mostly shared between the two genera, and built around the same physiological priorities in both. The passage between them is not a switch: the type III apparatus is dismantled completely, the Nod factor operons are attenuated but kept, and the *fixABCX* operon under a *nod* box is amplified, so that membership of the early programme is a property of the operon rather than of the regulatory element above it. Across three legumes of one strain, the variable that organises what differs is the identity of the host and not the type of nodule it forms, and host identity acts on whether a gene is mobilised and at what amplitude, almost never on the direction of its change.

## CRediT authorship contribution statement

S. Acosta-Jurado: Validation, Investigation, Methodology; F. Fuentes-Romero: Validation, Investigation, Methodology; P. Navarro-Gómez: Investigation; P. Ayala-García: Investigation; F.J. Ollero: Conceptualization; J.I. Jiménez-Zurdo: Project Administration, Funding Acquisition; F. Pérez-Montaño: Conceptualization, Writing - Original Draft Preparation; J.M. Vinardell: Funding Acquisition, Project Administration, Writing - Review & Editing; F.J. López-Baena: Funding Acquisition, Project Administration, Writing - Review & Editing.

## Declaration of competing interest

The authors declare that they have no known competing financial interests or personal relationships that could have appeared to influence the work reported in this paper.

## Declaration of generative AI and AI-assisted technologies in the writing process

The experimental design, the planning and execution of all experiments, the interpretation of the data and the logical structure and argument of this article are entirely the work of the authors. Claude Science (Anthropic) was used as a computational assistant for parts of the data analysis and for the preparation of figures and tables, and for language editing. The authors reviewed and verified all content, including every value reported, and take full responsibility for it.

## Acknowledgements

F. Fuentes-Romero was supported by the Junta de Andalucía and NextGenerationEU/PRTR through a predoctoral grant PREDOC_01119. P. Ayala-García received an FPU fellowship (FPU18/06248) from the Spanish State Research Agency (AEI) (MCIN/AEI/10.13039/501100011033). This work was funded by a research grant from the State Subprogram for Knowledge Generation of the Spanish Ministry of Science, Innovation and Universities (MICIU), the AEI and the European Union, reference PID2022-141156OB-I00, awarded to F.J. López-Baena and J.M. Vinardell, and by project P20_00185 from the Junta de Andalucía, awarded to J.I. Jiménez-Zurdo and J.M. Vinardell.

## Data availability

Raw sequence reads have been deposited in the NCBI Sequence Read Archive under BioProject accessions PRJNA1557975 (*Sinorhizobium fredii* HH103 transcriptome) and PRJNA1557978 (*Rhizobium tropici* CIAT 899 transcriptome).

## Appendix A. Supplementary data

Supplementary tables S1-S11 are provided as a single workbook, with a contents sheet (S0) summarising the criteria behind each table: S1 annotated DEGs by condition (11,025 rows), S2-S3 host-specificity partitions, S4 curated functional-block enrichment, S5 orthologous pairs, S6 validation against the 2016 dataset, S7 full annotation of both strains (13,392 features), S8 COG category enrichment, S9 per-gene COG assignment (10,235 proteins), S10 recovery of the 101 genistein-responsive ORFs of 2016 in HH103 and S11 recovery of the 20 qRT-PCR-validated genes of 2016 in CIAT 899.

## Notes

### Competing Interest Statement

The authors have declared no competing interest.

### Summary of Updates

This revised version does not alter any substantive findings or conclusions, but it refines and improves many details throughout the manuscript

